# ERVcaller: Identify polymorphic endogenous retrovirus (ERV) and other transposable element (TE) insertions using whole-genome sequencing data

**DOI:** 10.1101/332833

**Authors:** Xun Chen, Dawei Li

## Abstract

**Motivation:** Approximately 8% of the human genome is derived from endogenous retroviruses (ERVs). In recent years, an increasing number of human diseases have been found to be associated with ERVs. However, it remains challenging to accurately detect the full spectrum of polymorphic (unfixed) ERVs using next-generation sequencing (NGS) data.

**Results:** We designed a new tool, ERVcaller, to detect and genotype transposable element (TE) insertions, including ERVs, in the human genome. We evaluated ERVcaller using both simulated and real benchmark whole-genome sequencing (WGS) datasets. By comparing with existing tools, ERVcaller consistently obtained both the highest sensitivity and precision for detecting simulated ERV and other TE insertions derived from real polymorphic TE sequences. For the WGS data from the 1000 Genomes Project, ERVcaller detected the largest number of TE insertions per sample based on consensus TE loci. By analyzing the experimentally verified TE insertions, ERVcaller had 94.0% TE detection sensitivity and 96.6% genotyping accuracy. PCR and Sanger sequencing in a small sample set verified 86.7% of examined insertion statuses and 100% of examined genotypes. In conclusion, ERVcaller is capable of detecting and genotyping TE insertions using WGS data with both high sensitivity and precision. This tool can be applied broadly to other species.

**Availability:** www.uvm.edu/genomics/software/ERVcaller.html

**Supplementary information:** Supplementary data are available at *Bioinformatics* online.

## Introduction

### Origin of endogenous retroviruses

Transposable elements (TEs), which were first found in maize, are groups of mobile DNA sequences that collectively comprise a large percentage of most eukaryotic genomes, e.g., 44% of the human genome (Mills, et al., 2007). TEs can be categorized into two classes based on whether their transposition intermediate is RNA or DNA: Class I (RNA) retrotransposons, and Class II (DNA) DNA transposons (Wessler, 2006). There are four types of mobile elements in the human genome, including Alu, long interspersed element-1 (LINE1), short interspersed nuclear element/variable number tandem repeat/Alu (SVA), and endogenous retrovirus (ERV). ERVs are a unique group of Class I TEs resulting from the fixation of ancient retroviral infections and integrations into the host genome (Katzourakis, et al., 2007). In the human genome, 8% of all nucleotides are from ERVs (International Human Genome Sequencing, 2001; Jern and Coffin, 2008). The human ERVs (HERVs) as a whole are fairly old, dating back 35 million years. Most, if not all HERVs, are not replication-competent due to accumulated mutations, insertions, deletions, or having become solo-long terminal repeats (solo-LTRs) (Jern and Coffin, 2008; Kazazian and Moran, 2017; Marchi, et al., 2014). HERV K family (HERV-K) are relatively younger, having integrated in the past million years. Insertionally polymorphic HERV-K insertions have been reported in the human populations (Belshaw, et al., 2005; Kahyo, et al., 2017; Macfarlane and Badge, 2015; Wildschutte, et al., 2016).

### Endogenous retroviruses and human diseases

ERVs in the human genome have been shown to effect both the genome structure and biological functions (Kazazian and Moran, 2017; Navarro, 2017). ERVs can lead to genomic rearrangements through non-allelic homologous recombination with other ERV copies (Thomas, et al., 2018). ERVs can also behave as a source of promoters, enhancers, or transcriptional factor binding sites for regulating human gene expression levels (Fuentes, et al., 2018; Jern and Coffin, 2008). They can either systematically transcribe stage-specific RNAs in early embryonic development (Goke, et al., 2015; Robbez-Masson and Rowe, 2015) or express under certain disease conditions (Rooney, et al., 2015), or by interaction with certain infectious agents, e.g., human immunodeficiency virus and Epstein-Barr virus (Garrison, et al., 2007; Handsaker, et al., 2011; Leung, et al., 2018). Expressed viral genes, e.g., *gag* and *pol*, can interact with the human transcriptome or modulate the human immune system (Jern and Coffin, 2008). Expression of ERVs has thus been associated with autoimmune diseases (Brodziak, et al., 2012; Groger and Cynis, 2018; Marguerat, et al., 2004), mental disorders (Douville and Nath, 2014; Li, et al., 2015; Slokar and Hasler, 2015), cancers (Gonzalez-Cao, et al., 2016; Kassiotis, 2014), and many other human diseases.

Compared to monomorphic (fixed) ERVs, which are shared by all humans and usually ancient retroviral integration events, polymorphic (unfixed) ERVs are typically more recently integrated (Belshaw, et al., 2005). Fixed ERVs are more likely to have been co-opted to serve a function for the host organism. For example, these ERVs contain motifs for gene regulation (Chuong, et al., 2013; Fort, et al., 2014; Jern and Coffin, 2008), and co-option of ERVs is a driving force behind the evolution of the human immune system (Chuong, et al., 2016). By comparison, the polymorphic ERVs are more likely to retain biological functions as “viruses”. The speculated pathogenic mechanisms of polymorphic ERVs include disrupting the function of human genes at or near their integration sites, expressing viral proteins such as Env, or altering the adaptive and innate immune response (Moyes, et al., 2007). Although the pathogenicity of polymorphic ERVs is still poorly understood due to the complexity of ERV sequences, some polymorphic ERVs have been found to be associated with human diseases (Belshaw, et al., 2005; Karamitros, et al., 2018; Marchi, et al., 2014). For example, a polymorphic HERV-K insertion in the *RASGRF2* gene was recently found to be significantly associated with intravenous drug abuse (Karamitros, et al., 2018). This study focuses on identifying genome-wide polymorphic ERVs and other TEs.

### Existing tools for detecting polymorphic ERVs and other TEs

To determine the pathogenic effects of ERVs, it is necessary to first identify the full spectrum of polymorphic ERV insertions in the human genome. Similar to single nucleotide polymorphisms (SNPs), the same ERV insertions can be found in one or more individuals, leading to polymorphic ERV loci. Indeed, many polymorphic ERV loci have been recently discovered in the human genome (Kahyo, et al., 2017; Macfarlane and Simmonds, 2004; Marchi, et al., 2014; Wildschutte, et al., 2016). For example, Lee *et al*. discovered a total of 15 polymorphic ERV loci by screening 44 individuals (Lee, et al., 2012). By screening additional samples, Emanuele *et al*. found 17 loci, including two novel loci (Marchi, et al., 2014), with an average of six polymorphic ERV insertions per human genome. By analyzing a different set of samples, Wildschutte *et al*. found 19 new loci (Wildschutte, et al., 2016). These studies indicate that more polymorphic ERV loci likely exist in the human population, which have not yet been discovered.

With the wide-spread application of next-generation sequencing (NGS) technologies, some bioinformatics tools have been used to discover polymorphic TEs. VariationHunter (Hormozdiari, et al., 2009; Hormozdiari, et al., 2010) and Hydra (Quinlan, et al., 2010), which were originally designed for detecting general structural variations, were first adapted for detecting polymorphic TE insertions using whole-genome sequencing (WGS) data. However, due to the long fragment insertions and highly abundant repeated sequences, the detection of polymorphic TEs, including ERVs, is more difficult than the detection of SNPs, small insertions and deletions (Indels), or other structural variations. Specific tools were then developed, including TEA (Lee, et al., 2012), RetroSeq (Keane, et al., 2013), TIF (Nakagome, et al., 2014), Mobster (Thung, et al., 2014), Tangram (Wu, et al., 2014), TEMP (Zhuang, et al., 2014), ITIS (Jiang, et al., 2015), STEAK (Santander, et al., 2017), and MELT (Gardner, et al., 2017) (**Supplementary Table 1**). Although each software has its own merits, many of these tools are limited or insufficient for accurately detecting or genotyping the full spectrum of TE insertions, particularly for ERVs. For example, most tools were designed to screen the consensus TE references only, and thus would potentially miss ERV and other TE insertions containing novel TE sequences or those having many point mutations or indels (Goerner-Potvin and Bourque, 2018).

### A new software: ERVcaller

In this study, we developed a novel software, ERVcaller, to detect and genotype genome-wide TE insertions, particularly ERVs, with paired-end or single-end WGS data. Due to the optimization of multiple operations, ERVcaller achieved both high sensitivity and precision, and thus, identified new ERV and other TE insertions, which could not be detected by most of the existing tools. ERVcaller generates the genotypes of the identified ERVs and other TEs in VCF files for subsequent analyses.

## Methods

### ERVcaller software algorithm

#### Extract unmapped and anomalous reads

ERVcaller first aligns raw FASTQ reads to the human reference genome using BWA-MEM (Li, 2013) with the default parameters. It can also directly use a pre-aligned BAM file generated by either BWA-MEM or other aligners as input. ERVcaller supports the use of multiple BAM files as inputs for a single sample. ERVcaller then extracts reads that either do not align or align anomalously to the human genome using Samtools (Li, et al., 2009). Specifically, to obtain one-end unmapped reads, ERVcaller extracts (selects) reads flagged as unmapped (SAM flag=4) and then removes those also flagged as mate unmapped and non-primary alignment (SAM flag=264). To obtain one-end mapped reads, ERVcaller extracts reads flagged as mate unmapped (SAM flag=8) and then removes those also flagged as read unmapped and non-primary alignment (SAM flag=260). To obtain anomalous reads, ERVcaller extracts reads flagged as neither mapped in a proper pair nor having both the read and its mate-pair unmapped (SAM flag≠14) and then extracts the reads with one end uniquely aligned to the human reference genome and the other end aligned to multiple locations. To obtain soft-clipped reads, ERVcaller extracts reads flagged as being mapped in a proper pair (SAM flag=2) and then extracts the unmapped portions of soft-clip reads (≥ 20 base pair (bp)) in FASTQ format using SE-MEI (https://github.com/dpryan79/SE-MEI) (**Supplementary Figure 1**).

#### Align the obtained reads against the TE references

ERVcaller aligns all obtained reads concurrently against all TE references using BWA-MEM with customized parameters (**Supplementary Table 2**). Three types of supporting reads are extracted for detecting TE insertion events. Chimeric reads (also known as discordant reads) are defined as a read pair having one end aligned to the human reference genome while the other end does not align to the human reference genome but aligns to a TE reference sequence(s) (Chen, et al., 2018). Improper reads are defined as a read pair having both ends aligned to the human reference genome but not the same chromosome or within the insert size of each other, and one end uniquely aligned to the human reference and the other end aligned to a TE sequence. Split reads (also known as soft-clipped reads) are reads with one portion aligning to the human reference genome (the two ends still mapped in a proper pair) and the other portion aligned to a TE reference sequence(s) (Chen, et al., 2018) (**Supplementary Figure 2**).

#### Detect genome-wide ERV and other TE insertions and perform quality controls

ERVcaller first aligns all obtained supporting reads against the human reference genome, and then groups the reads in windows of two insert-sizes to identify the potential genomic regions containing TE insertions. ERVcaller then determines the TE insertion candidates based on the number of reads and the average alignment scores of supporting reads within each genomic region. As the human genome contains many fixed TE sequences, which have high sequence similarity to polymorphic TEs, ERVcaller aligns the supporting reads of each polymorphic TE candidate to the human reference sequence within each candidate genomic region. The TE candidates with no confident supporting reads are removed, i.e., when the supporting reads fully align to the human reference sequence of the candidate region.

#### Fine map insertion breakpoints at single nucleotide resolution

Once a TE insertion event is detected, ERVcaller uses the chimeric, improper, and split reads to determine the chromosomal location of each breakpoint (junction) at nucleotide resolution. If split reads spanning the breakpoints are detected, the breakpoint locations can be precisely determined. Without the presence of split reads, ERVcaller uses the chimeric and improper reads, including those crossing the breakpoints, to estimate the breakpoint location. Specifically, if either the upstream (5’) or downstream (3’) breakpoint of the insertion is identified, the first nucleotide of the nearest read to the breakpoint is considered to be the estimated breakpoint; if both the upstream and downstream breakpoints are identified, the median of the first nucleotide of the nearest read to the breakpoints is considered to be the estimated breakpoint.

#### Genotype ERV and other TE insertions and output as VCF files

The total number of chimeric, improper, and split reads supporting the hypothesis of the polymorphic TE insertion versus the total number of reads supporting the hypothesis of no insertion is used to determine the genotype of the TE insertion. The reads supporting no insertion are those which can be fully aligned to the human reference genome across the breakpoint. ERVcaller extracts these reads from the BAM file using Samtools and in-house Perl scripts. If the number of reads supporting the presence of a polymorphic TE insertion and the number of reads supporting no insertion at a locus are the same, the genotype of this TE insertion is defined to be heterozygous (**Supplementary Figure 3**). However, genotyping is often complicated by noise derived from the sequencing process, alignment, and subsequent analyses. For example, for a heterozygous TE insertion, the number of reads supporting the presence of a TE insertion is likely to be smaller than that supporting the absence of the insertion due to the experimental and bioinformatics challenges in capturing low complexity sequences. Thus, by default, ERVcaller outputs all the TE insertions that contain a minimum of three supporting reads, including at least one chimeric or improper read; however, the TE insertions should be further filtered using appropriate criteria relative to the total number of reads or matched with the sequencing depth, e.g., a larger number of supporting reads is required for high-depth data. Consequently, ERVcaller calculates the genotype likelihood to determine the homozygous versus heterozygous status of each identified polymorphic TE insertion using a combined insert-size and read-depth approach (Handsaker, et al., 2011). Finally, ERVcaller outputs the genotypes of genome-wide ERV and other TE insertions in Variant Call Format (VCF) (Danecek, et al., 2011), which may be used for subsequent genetic association analysis. With the implemented scripts, the outputs from multiple samples can also be either matched to a list of previously reported consensus TE loci or to the TE loci detected in those samples.

### Software comparison

To compare ERVcaller with the existing tools for detecting ERV and other TE insertions, we selected five of the most recently published tools, including ITIS (Jiang, et al., 2015), TEMP (Zhuang, et al., 2014), RetroSeq (Keane, et al., 2013), STEAK (Santander, et al., 2017), and MELT (Gardner, et al., 2017). After compilation and installation, we ran each tool with the default parameters. We used BWA-MEM to align raw reads to the human reference genome, and then used Samtools to convert the generated SAM files to indexed BAM files. Aside from ITIS, which uses FASTQ files as input, ERVcaller and the other four tools used BAM files as input. An additional TE annotation file was provided for MELT, TEMP, and RetroSeq, as required or recommended by these tools. MELT-SINGLE was applied to detect TE insertions. Only the TE insertions (events) with three or more supporting reads were kept for software comparison. To examine the runtimes of these tools, we used 12 core computing nodes with 48 GB memory from the Vermont Advanced Computing Core cluster.

### Reference sequences for detecting ERV and other TE insertions

Four TE consensus sequences, including one Alu, one LINE1, one SVA, and one HERV-K, were used for each tool to ensure equal comparisons. The human reference genome build GRCh38 (hg38) downloaded from the UCSC Genome Browser was used. To examine whether the use of different TE references influences the performance of ERVcaller, we compared the results obtained by screening three different TE reference libraries, including (1) the four TE consensus references, (2) the 23 TE consensus references obtained from Tangram (Wu, et al., 2014), including 17 LINE1, four Alu, one HERV-K, and one SVA sequences, and (3) the entire Repbase database containing all eukaryotic repetitive DNA elements (Bao, et al., 2015).

### Simulation

Due to the biological importance of ERVs, we first simulated ERV insertions using an in-house Perl script, and then examined the sensitivity and precision of ERVcaller and the five existing tools for detecting polymorphic ERV insertions. Specifically, we randomly inserted modified full length and partial HERV-K consensus sequences into the human chromosome 1 accessible locations that were masked by the 1000 Genomes Project (The Genomes Project, 2015) (referred to as “modified full consensus HERV-K insertions” and “modified partial consensus HERV-K insertions” below, respectively). The HERV-K consensus sequences were modified based on known features, such as orientations and mutations. The genomic locations of the simulated ERV insertions were matched with the distribution of real polymorphic ERV insertions reported in the human genome (Chaisson, et al., 2018; Wildschutte, et al., 2016). Paired-end reads of 100 bp length and a 500 bp insert size were simulated using pIRS (Hu, et al., 2012) with the default parameters. Two series of data were simulated, including 500 full length and 500 partial (randomly fragmented) HERV-K insertions.

To evaluate other TE insertions, we generated two additional series of data containing all TE types, including Alu, LINE1, SVA, and HERV-K. TE sequences were randomly inserted into the human genome according to the number and distribution of each TE type in the 1000 Genomes Project benchmark sample NA12878, i.e., 921 Alu, 124 LINE1, 46 SVA, and 7 HERV-K (Gardner, et al., 2017; Sudmant, et al., 2015; Wildschutte, et al., 2016). The first series contained TE consensus sequences that we randomly modified based on the known TE features, including length, orientation, mutations, 3’ transduction, and target site duplications (referred to as “modified TE consensus insertions” below). The second series contained TE insertions derived from real polymorphic TE sequences that we obtained from the dbRIP database (Wang, et al., 2006). Each series was simulated at multiple sequencing depths, including 1X, 5X, 10X, 15X, 20X, 25X, 30X, and 60X.

### 1000 Genomes Project samples and data preprocessing

To evaluate the performance of ERVcaller with real WGS data, we applied ERVcaller, MELT, RetroSeq, and TEMP to the low-coverage (~8.6X depth) WGS data of 45 1000 Genomes Project samples (Genomes Project, et al., 2010) (**Supplementary Table 3**). CRAM files (hg38) downloaded from the International Genome Sample Resource database were converted using Samtools to indexed BAM files, which were then used as input for each tool. The same parameters used for the simulated datasets were used for this real data. To compare the performance of these tools, we matched the detected TE insertions to 21,723 consensus TE loci. Consensus TE loci were defined to be the TE loci (locations) that have been reported previously in multiple samples. To generate these consensus loci, we collected a total of 19,420 TE loci based on the TE insertions maintained in the dbRIP database (Wang, et al., 2006), and those identified in 2,535 samples from the 1000 Genomes Project (Sudmant, et al., 2015; Wildschutte, et al., 2016), e.g., ~80% of the TE loci in the 1000 Genomes Project samples occurred in two or more samples. Additionally, we also considered a small set (3,263) of candidate TEs selected based on the complementary data from no more than nine samples (Chaisson, et al., 2018) (**Supplementary Table 4**). We then removed redundant TE loci by merging all these TE loci using a 500 bp window, resulting in a total of 21,723 consensus TE loci.

We also analyzed the high-coverage (~68.5X depth) WGS data of three 1000 Genomes project samples with experimentally validated Alu, LINE1, and SVA insertions (**Supplementary Table 3**). These insertions were detected using Spanner (https://github.com/chipstewart/Spanner) and validated by polymerase chain reaction (PCR) (Stewart, et al., 2011). We downloaded the raw FASTQ files from the International Genome Sample Resource database and aligned them to the human reference genome using BWA-MEM with the default parameters. The resulting SAM files were converted to sorted and indexed BAM files using Samtools, which were then used as inputs for each tool. As the genome build hg18 was used in the original paper (Stewart, et al., 2011), the hgLiftOver tool (https://genome.ucsc.edu/cgi-bin/hgLiftOver) was used to convert the coordinates to hg38.

### PCR and Sanger sequencing validation

To evaluate the detection and genotyping accuracies of ERVcaller, we performed molecular experiments to verify the insertion status and genotypes of ERVs and other TEs identified by ERVcaller in a total of six samples (i.e., the 1000 Genomes Project samples), for which DNA specimens were available to our laboratory. We first selected two ERV insertion events that had not been previously experimentally verified and performed PCR to verify the loci in two samples. The primers for the two loci have been described previously (Wildschutte, et al., 2016). The PCR products were purified and cloned using the StrataClone Blunt PCR Cloning Vector pSC-B-amp/kan plasmid (Agilent Technologies, La Jolla, CA). We then used Sanger sequencing to verify the ERV genotypes, i.e., the alleles with and without the insertion. Based on the Sanger sequencing results, we then used PCR to verify the genotypes of the two loci in the remaining samples. For other TEs, we used PCR to verify the insertion status of 15 Alu insertions that were newly identified in these samples by ERVcaller. The primers for PCR were designed based on the 300 bp upstream and downstream sequences using Primer3 with default parameters (Untergasser, et al., 2012) (**Supplementary Table 5**).

## Results

### ERVcaller pipeline

We designed a new bioinformatics tool, ERVcaller, for the accurate detection and genotyping of polymorphic ERV and other TE insertions. ERVcaller is composed of three modules, including **a**) extracting unmapped and anomalous reads, **b**) obtaining supporting reads, and **c**) detecting and genotyping TE insertions (**Figure 1**). The inputs for ERVcaller include either FASTQ or BAM file(s), the human reference genome, and TE reference sequence(s). ERVcaller first extracts all reads that could not be fully mapped to the human reference genome and obtains the anomalous reads in which the two ends are not aligned in a proper pair. These reads are then aligned to the TE reference library. ERVcaller uses three types of supporting reads, i.e., chimeric, split, and improper reads, to determine the chromosomal location of each TE insertion, including both upstream and downstream breakpoints when possible (**Supplementary Figure 2**). ERVcaller outputs every TE insertion meeting the following three criteria: 1) It has at least three supporting reads (by default), at least one of which is a chimeric or improper read; 2) the average alignment score of each supporting read is greater than 30; and 3) each read contains at least 50 bp of sequence mapped to the human reference genome. If no reads could be uniquely mapped to the human genome, the TE insertion would likely be a false positive; however, as some ERV insertions have been found within repeat regions, i.e., segmental duplications and regions containing fixed TEs (Wildschutte, et al., 2016), ERVcaller annotates and retains these low confidence TEs in a separate output file for further manual investigation. After stringent filtering, high confidence TE insertions are then genotyped based on the number of reads crossing the breakpoints, as described in the Methods section (**Supplementary Figure 3**). ERVcaller saves the resulting genome-wide ERVs and other TEs in VCF files.

**Figure 1.**
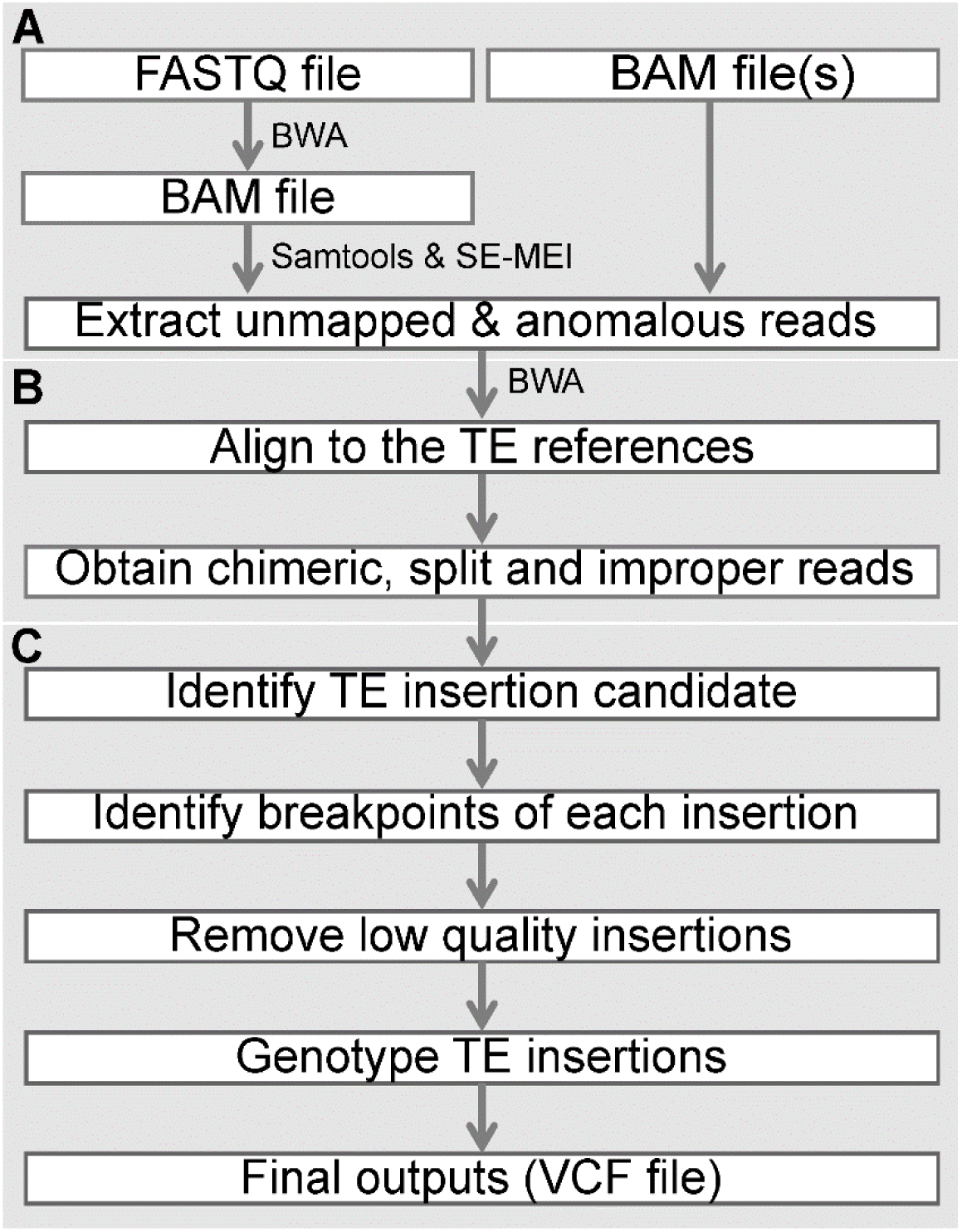
ERVcaller workflow. The three major components include **A**) extracting unmapped and anomalous reads; **B**) obtaining supporting reads; and **C**) detecting and genotyping ERV and other TE insertions.

### Comparative analyses of detection of simulated ERV and other TE insertions

We performed a comprehensive comparison for detecting ERV and other TE insertions between ERVcaller and five existing tools, including RetroSeq, STEAK, ITIS, TEMP, and MELT (**Supplementary Table 1**). To measure sensitivity and precision, we used the TE consensus sequences as the reference and applied each tool to a total of four series of simulated data containing, 1) modified full consensus HERV-K insertions, 2) modified partial consensus HERV-K insertions, 3) modified consensus TE insertions, and 4) TE insertions derived from real polymorphic TE sequences. ERV and other TE types were evaluated separately. Sensitivity (detection power) was defined as the ratio of correctly identified versus total simulated TE insertions. Precision was defined as the ratio of correctly identified versus total identified TE insertions. The sensitivity and precision were calculated at sequencing depths from 1X to 60X. Additionally, the accuracies for detecting insertion breakpoints and the runtimes were compared among all six tools.

We first compared the runtimes of the six tools. The four series of simulated data were combined for this analysis. ERVcaller had the shortest runtime (6.9 mins) at a sequencing depth of 30X with 12 cores and 48 GB memory, followed by TEMP (8.8 mins), RetroSeq (16.6 mins), STEAK (51.5 mins), MELT (63.5 mins), and ITIS (553.1 mins) (**Figure 2A**). We further evaluated the runtime of ERVcaller for three TE libraries, including the four TE consensus references, the 23 consensus references, and the entire Repbase database. ERVcaller only had a slightly longer runtime (8.4 mins) when using the entire Repbase database (**Figure 2B**). This high speed allows ERVcaller to identify the full spectrum of ERV and other TE insertions in a large number of samples.

**Figure 2.**
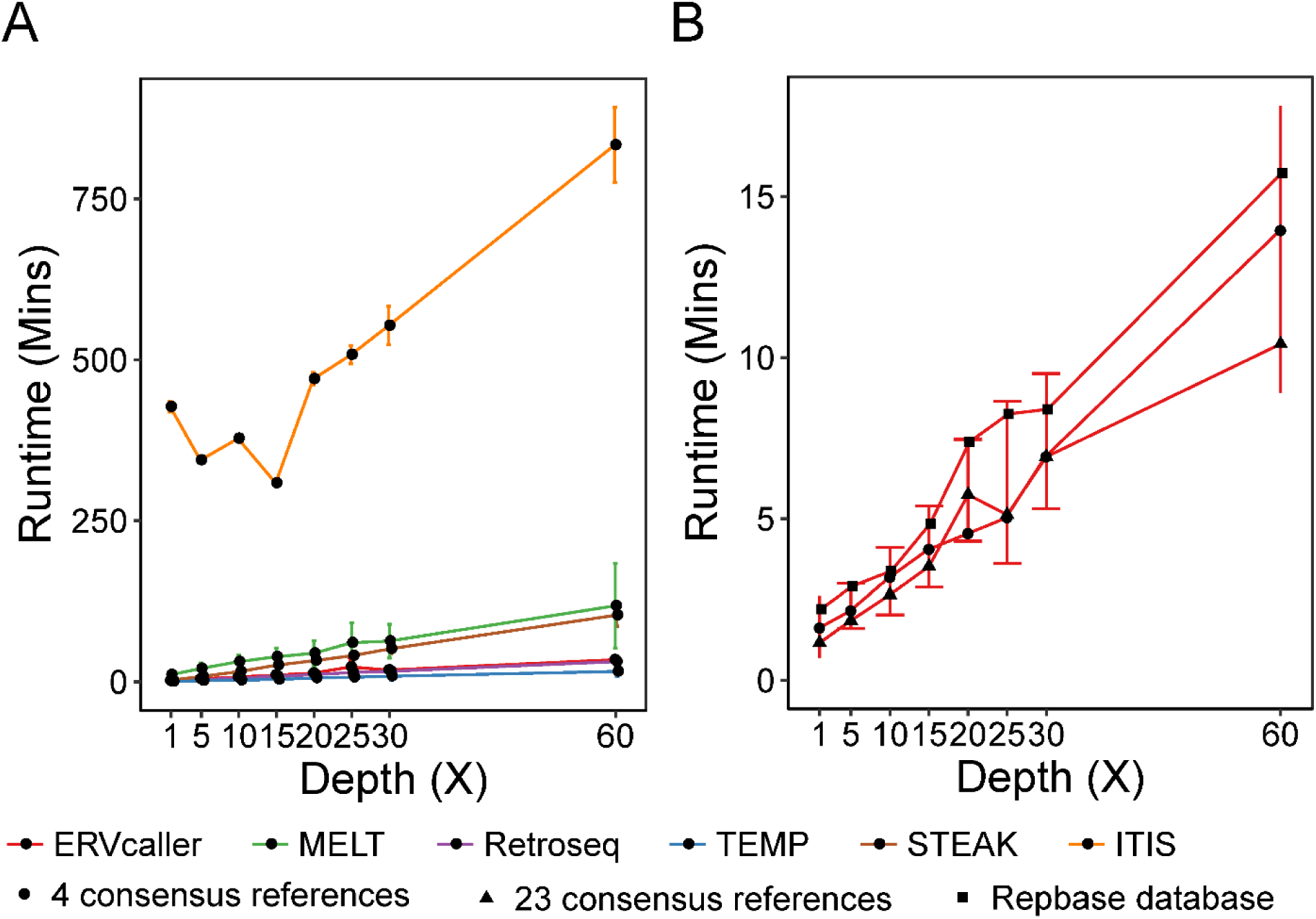
Comparison of runtimes of ERVcaller with the five existing tools for analyzing the simulated datasets. **A**) Four consensus references were used by each tool. **B**) Three reference libraries were used by ERVcaller, including the four consensus references, the 23 consensus references, and the Repbase database.

We then measured the sensitivity and precision of the six tools for detecting HERV-K insertions. For detecting full HERV-K insertions, all six tools achieved a high sensitivity (> 95%) when the sequencing depth was 30X or higher (**Figure 3A**). ERVcaller, RetroSeq, and MELT achieved > 98% precision, while TEMP, STEAK, and ITIS had 88%, 26%, and 13% precision, respectively (**Figure 3A**). For detecting the partial HERV-K insertions, ERVcaller, RetroSeq, and TEMP consistently achieved high sensitivity (> 95%) at 30X depth, while ITIS and STEAK only had < 20% sensitivity (**Figure 3B**). Aside from ITIS and STEAK (< 10%), all four other tools showed high precision (> 95%) (**Figure 3B**). Thus, for detecting the full and partial HERV-K insertions, only ERVcaller, RetroSeq, and MELT achieved both high sensitivity and precision among the six tools analyzed.

**Figure 3.**
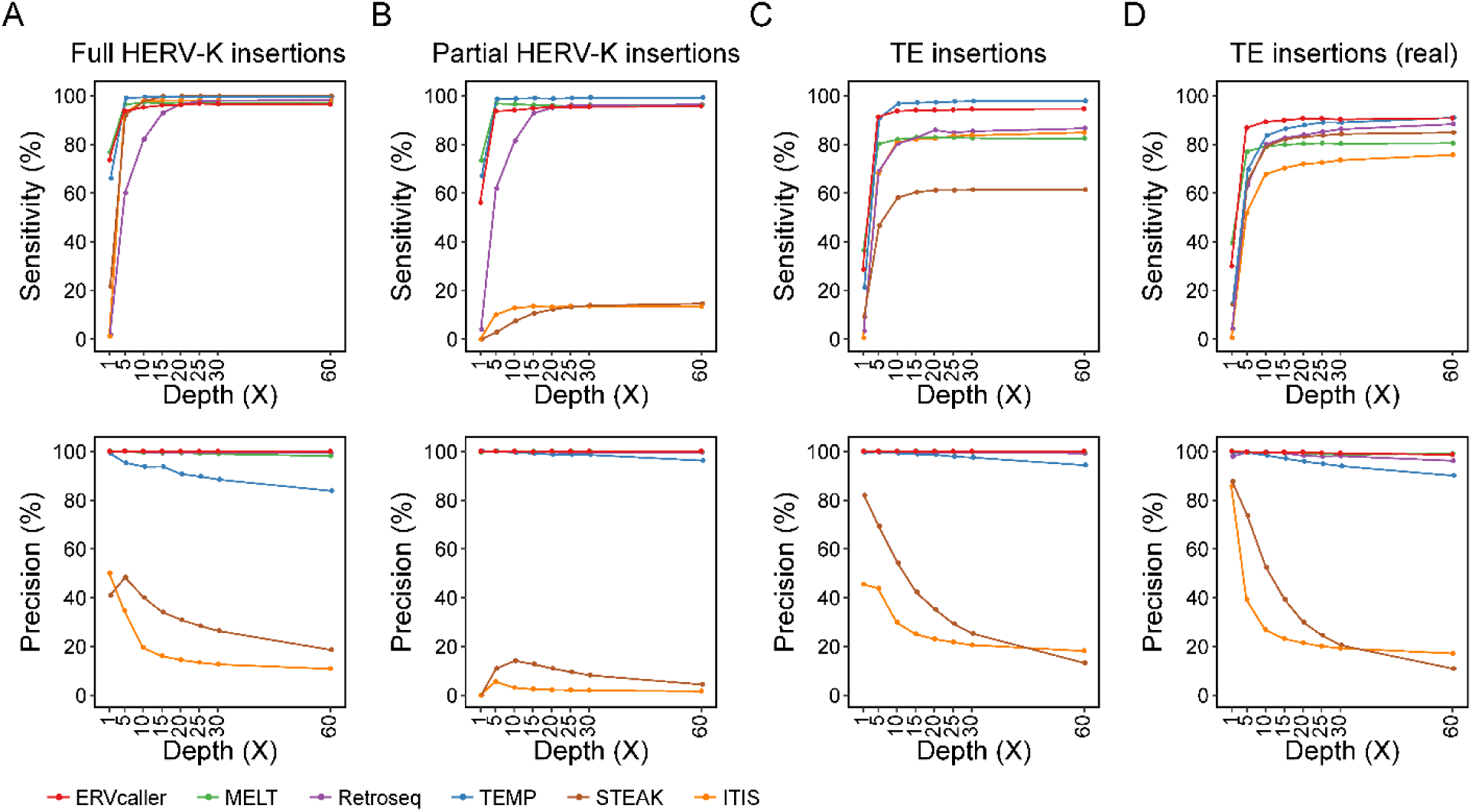
Comparison of sensitivity and precision of ERVcaller with the five existing tools for analyzing four different simulated datasets, including **A**) the dataset with modified full consensus HERV-K insertions; **B**) the dataset with modified partial consensus HERV-K insertions; **C**) the dataset with modified consensus TE insertions; and **D**) the dataset with TE insertions derived from real polymorphic TE sequences.

We further evaluated the performance of the tools for detecting all TE insertions, including Alu, LINE1, SVA, and HERV-K. For detecting the modified consensus TE insertions, only TEMP (98%) and ERVcaller (94%) had consistently high sensitivity at 30X depth (**Figure 3C**), while RetroSeq, ITIS, and MELT had a decreased sensitivity (~84%), and STEAK had the lowest sensitivity (61%) (**Supplementary Figure 4**). ERVcaller, RetroSeq, TEMP, and MELT consistently had > 98% precision, while STEAK and ITIS had < 25% precision (**Figure 3C**). For detecting the TE insertions derived from real polymorphic TE sequences, ERVcaller achieved the highest sensitivity (90%) at 30X, followed by TEMP (89%), RetroSeq (86%), STEAK (84%), MELT (80%), and ITIS (73%) (**Figure 3D**). The precision remained similar, i.e., > 98% for ERVcaller, RetroSeq, TEMP, and MELT, and < 25% for STEAK, and ITIS (**Figure 3D**). All six tools had decreased sensitivity when analyzing the real polymorphic TE insertions. However, ERVcaller was the least affected compared to the five existing tools, retaining both the highest sensitivity and precision among the six tools.

To compare the accuracies for detecting breakpoints, we only evaluated four of the six tools because ITIS and STEAK were not designed to detect breakpoints at single nucleotide resolution. If the detected breakpoint coordinates were within a 2-bp window of the simulated coordinate, it was considered correctly detected. We found that MELT and ERVcaller showed consistently higher accuracy, e.g., > 90% at depths ≥ 30X, than RetroSeq and TEMP. The latter two obtained significantly lower accuracies, particularly for detecting TE insertions from the consensus sequences and the dbRIP database (**Supplementary Table 6**). Thus, both MELT and ERVcaller are capable of correctly detecting the breakpoints of ERVs and other TEs, regardless of the insertion length or TE type.

To evaluate whether repetitive and redundant sequences in the TE references influenced the sensitivity and precision of ERVcaller, we analyzed all the four simulated datasets by screening three different TE libraries. When the four consensus references were used, ERVcaller consistently showed high sensitivity (> 95% and > 88%, respectively) and precision (> 99%) for detecting the HERV-K and other TE insertions at depths ≥ 30X (**Supplementary Figure 5**). When the 23 consensus references were used, ERVcaller showed the same sensitivity and precision as for the four consensus references. When the whole Repbase database was used, the sensitivity only decreased 2 – 5%, and the precision remained > 99%. Hence, ERVcaller achieved both high sensitivity and precision without being significantly influenced by the selection of TE references. This suggests that larger TE reference libraries, presumably containing additional ERVs and other TEs, may be valuable for ERVcaller to detect novel or highly mutated ERV and other TE insertions.

### Comparative analyses of detection of ERV and other TE insertions in 1000 Genomes Project samples

To further evaluate the performance of ERVcaller in analyzing real benchmark data, we analyzed the WGS data of 45 samples from the 1000 Genomes Project. Due to the low precision of ITIS and STEAK shown in the above *in silico* analyses, we compared ERVcaller with only three other tools, including MELT, RetroSeq, and TEMP. We first compared each detected TE locus with the 21,723 consensus TE loci (locations) that we curated based on a large number of existing datasets (**Supplementary Table 4**) (described in the Method section). A TE insertion was considered to be correctly detected if its coordinate was matched within a 500 bp window of one of these consensus TE loci. ERVcaller detected 7,607 consensus TE loci in the 45 samples, including 5,945 Alu, 902 LINE1, 311 SVA, 19 HERV-K, and 430 complex or mixed TE loci. By comparison, MELT, RetroSeq, and TEMP detected 7,189, 7,845, and 5,346 TE loci, respectively. A total of 4,121 were detected by all the four tools. The TE insertions detected by two or more tools were considered more confident than those detected by only one tool. ERVcaller detected the largest number of consensus TE loci (7,371) that were also identified by at least one other tool; by comparison, MELT detected 6,844 loci, followed by RetroSeq (6,808), and TEMP (5,130). On the other hand, only 488 out of the 21,723 loci were detected by at least two other tools but not detected by ERVcaller, while the corresponding numbers were 1,015, 1,053, and 2,647 for MELT, RetroSeq, and TEMP, respectively. Thus, ERVcaller detected the largest number of consensus (i.e., previously published) TE loci compared to the other three tools regarding detection of consensus ERV and other TE loci (**Figure 4A**).

**Figure 4.**
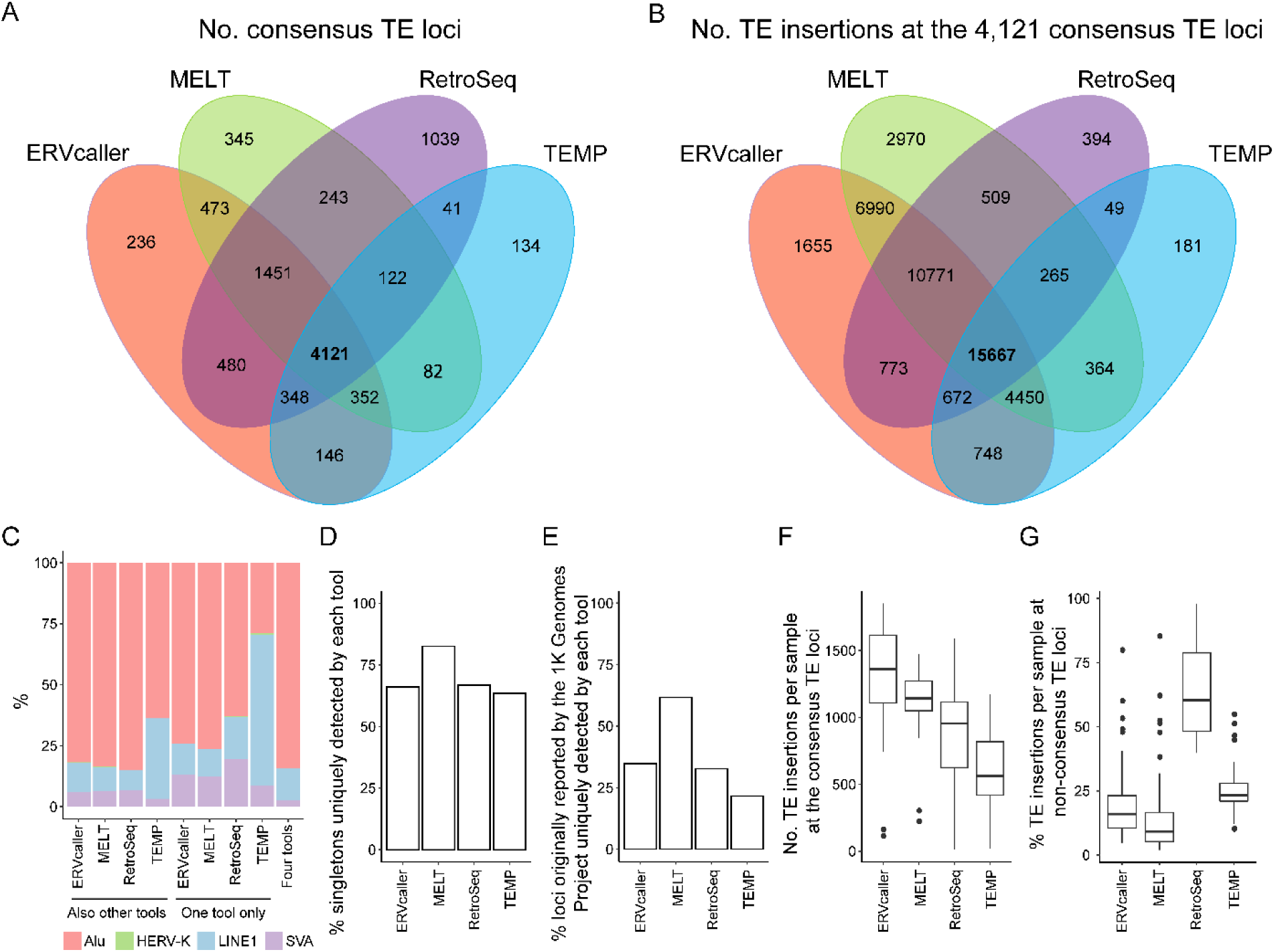
Comparison of ERVcaller with three other tools for analyzing the 45 samples from the 1000 Genomes Project. **A**) Comparison of the numbers of consensus TE loci identified in the 45 samples. **B**) Comparison of the total numbers of TE insertions at the 4,121 consensus TE loci identified in the 45 samples. **C**) The proportions of TE types for different tools. **D**) The percentages of singleton consensus TE loci detected by each tool only. **E**) The percentages of the consensus TE loci detected by each tool alone based on data reported by the 1000 Genomes Project (Sudmant, et al., 2015). **F**) The average numbers of TE insertions per sample at the consensus TE loci detected by each tool. **G**) The percentages of TE insertions per sample that were not from the consensus TE loci detected by each tool.

We then analyzed the total number of TE insertions (events) identified at the 4,121 consensus TE loci that were detected by all the four tools in the 45 samples (**Figure 4B**). MELT (41,986) and ERVcaller (41,726) detected comparable numbers of TE insertions at those loci. RetroSeq and TEMP only detected 29,100 and 22,396 TE insertions respectively. Furthermore, ERVcaller and MELT jointly detected a total of 37,878 TE insertions, accounting for 90% of the TE insertions detected by each. Thus, ERVcaller and MELT both detected significantly more TE insertions at the consensus TE loci than RetroSeq and TEMP.

We analyzed the proportion of each TE type for the TE loci that were detected by each tool alone. We found that ~60% of the TE insertions only detected by TEMP were LINE1 (**Figure 4C**). By comparison, LINE1 insertions detected by the other tools consistently accounted for < 20% of the total TEs. Indeed, in the benchmark NA12878 sample, the proportions of Alu, LINE1, SVA, and HERV-K were 83.9%, 11.3%, 4.19%, and 0.6%, respectively (Sudmant, et al., 2015; Wildschutte, et al., 2016). We then analyzed the percentage of singleton TE insertions (i.e., those observed in only one of the 45 samples) among the TE loci that were detected by each tool alone. We found that 83% of the TE loci only detected by MELT were singletons, while ERVcaller and the two other tools consistently had a percentage of ~65% (**Figure 4D**). Approximately, 62% of the TE loci only detected by MELT were already reported by the 1000 Genomes Project (Sudmant, et al., 2015), while ERVcaller and the two other tools consistently had < 35% (**Figure 4E**). This might be partially explained by the fact that MELT was originally developed for the 1000 Genomes Project (Sudmant, et al., 2015) and was used to generate most of the 21,723 consensus TE loci that we used in this paper.

We also analyzed the average number of TEs detected per sample by each tool (**Supplementary Table 7**). ERVcaller detected a total of 1,311 TE insertions at the consensus TE loci, followed by MELT (1,133), RetroSeq (893), and TEMP (607) (**Figure 4F**). Thus, ERVcaller detected 13.6%, 31.9%, and 53.7% more TE insertions per sample compared to MELT, RetroSeq, and TEMP, respectively. On the other hand, we analyzed the percentage of TE insertions that were not from the consensus TE loci. We found that an average of 21% of the TE insertions detected by ERVcaller were not from the consensus TE loci, while the average percentages for MELT, TEMP, and RetroSeq were 16%, 25%, and 65%, respectively (**Figure 4G**). The lower percentage (16%) achieved by MELT might also be explained, at least partially, by the reason outlined above. Thus, compared to the other tools, ERVcaller detected the largest number of TE insertions at the consensus TE loci.

We additionally analyzed the polymorphic HERV-K insertions detected in the 45 samples. ERVcaller identified a total of 281 HERV-K insertions in these samples. By comparison, TEMP, MELT, and RetroSeq detected 315, 912, and 1,118, respectively. When we matched the HERV-K insertions to the consensus HERV-K loci, most of which have been experimentally verified (Wildschutte, et al., 2016), we found that 94.7% of all HERV-K insertions detected by ERVcaller were matched with the consensus HERV-K loci. By comparison, the percentages for TEMP, MELT, and RetroSeq were 63.8%, 39.1%, and 24.7%, respectively. Thus, ERVcaller detected the highest percentage of confident HERV-K insertions among the consensus (mostly experiment-verified) HERV-K loci.

Based on the 45 samples, ERVcaller detected an average of 6.2 HERV-K insertions per individual, ranging from 1 to 10, involving a total of 30 different loci (**Supplementary Table 8**). Eight of the 30 loci have not been previously reported. Six of the eight loci were replicated by two or three of the evaluated tools, and the other two were replicated by either TEMP or RetroSeq. This result further indicates that ERVcaller can accurately detect new HERV-K insertions.

### Detection of experimentally verified TE insertions and genotypes

To measure the sensitivity of ERVcaller for detecting experimentally verified TE insertions, we first analyzed the WGS data of a trio from the 1000 Genomes Project, which contained PCR-verified insertion statuses for 1,136 TE insertions (events), including 1,029 Alu, 99 LINE1, and eight SVA (Stewart, et al., 2011). ERVcaller detected 97%, 87%, and 87.5% of them, respectively, or 95.7% when the three TE types were combined (**Supplementary Table 9**). Here, the ratio of correctly identified (experimentally verified) versus total examined (experimentally verified) TE insertions was defined to be the detection sensitivity. By comparison, MELT, RetroSeq, and TEMP obtained 97.3%, 96.9%, and 66.1% sensitivity, respectively, when the three TE types were combined. To further compare the sensitivity of ERVcaller with MELT, RetroSeq, and TEMP, we then analyzed another dataset. It included 28 Alu, 29 LINE1, and 25 SVA insertions that were previously reported by MELT in 21 samples from the 1000 Genomes Project (Gardner, et al., 2017), as well as six HERV-K insertions that were previously reported by RetroSeq in four additional samples (Keane, et al., 2013; Wildschutte, et al., 2016). All 88 insertions in the 25 samples had already been verified by PCR or Sanger sequencing. Our analysis showed that ERVcaller and MELT had comparable sensitivity (> 70%) in identifying these verified TE insertions, while RetroSeq and TEMP showed significantly lower sensitivity (< 45%) (**Supplementary Table 10**). Among the 88 insertions, 82 were originally reported by MELT; however, in our analysis, we only detected 60 of the 82 insertions using MELT, which might be due to our use of MELT parameters different from those used in its original paper.

To measure the accuracy of ERVcaller for genotyping TE insertions, we analyzed the same trio, which also contained PCR-verified genotypes for 1,080 TE insertions, including 970 Alu, 98 LINE1, and 12 SVA (Stewart, et al., 2011). ERVcaller correctly genotyped 98.0%, 83.3%, and 83.3% of them, respectively, or 96.6% when the three TE types were combined. We also analyzed the same data using MELT; the only other tool evaluated which was also able to generate TE genotypes. MELT correctly genotyped 95.2%, 84.7%, and 83.3% of them, respectively, or 94.1% when they were combined, slightly lower than ERVcaller (**Supplementary Table 9**).

### PCR and Sanger sequencing validation of new TE insertions and genotypes

As ERVcaller identified a total of 25,875 TE insertions that have never been previously reported in these 45 samples, we performed PCR for a total of 15 TE insertions that were newly identified by ERVcaller in the six samples with DNA specimens available to our laboratory. Thirteen out of the 15 TEs were verified. Among the 13 verified TEs, eight were identified by ERVcaller but not detected by any of the three other tools; three were able to be detected by MELT; and two were able to be detected by two or three other tools (**Supplementary Table 5**). The results indicate that ERVcaller is capable of detecting new TE insertions that might not be detected by other tools.

To evaluate the genotyping accuracy of ERVcaller, we chose two heterozygous HERV-K insertions that were identified by ERVcaller but had not been verified in two of the six samples with available DNA specimens. We first used Sanger sequencing to validate the genotypes of the two loci in the two samples (**Supplementary Figure 6**), and then used PCR to genotype the two loci in all six samples. All 12 genotypes detected by ERVcaller were verified in the six samples. One genotype was found to have been incorrectly determined by MELT (**Supplementary Table 11**).

## Discussion

Polymorphic ERV and other TE insertions are important categories of structural variations that are critical in human evolution and the development of human diseases (Burns, 2017; Jern and Coffin, 2008). The sequences of these variations are typically repetitive, and thus difficult to detect using short NGS reads. In this study, we improved upon the existing TE calling algorithms and developed ERVcaller to detect and genotype genome-wide ERV and other TE insertions using NGS data. By applying it to series of simulated datasets, we demonstrated that ERVcaller obtained both the highest sensitivity and precision when analyzing WGS data with a sequencing depth of ≥ 30X, except TEMP, which had higher sensitivity but lower precision. This was especially the case with the TE datasets simulated based on real polymorphic TE sequences, i.e., those in the dbRIP database. By further applying ERVcaller to 45 samples from the 1000 Genomes project, we detected the largest number of consensus TE loci that were consistently detected by at least one of the other tools that we evaluated. ERVcaller also identified the largest number of TE insertions per sample at these consensus TE loci. Based on our analyses of previously PCR-verified TE insertions, ERVcaller achieved high sensitivity (94.0%) for detecting TE insertions and high genotyping accuracy (96.6%). Our PCR and Sanger sequencing results in an additional small sample set also verified 86.7% (13 out of 15) of identified TEs and 100% of genotypes (i.e., the ratio of verified versus predicted genotypes, or 12 out of 12). ERVcaller also identified many new TE insertions that were not detected by any of the other tools.

ERVcaller achieves both higher sensitivity and precision than other tools for several reasons. ERVcaller implements a high mapping-rate aligner, BWA-MEM, and adopts customized parameters to align each read simultaneously to multiple references or locations. ERVcaller uses three types of supporting reads (chimeric, improper, and split reads), and determines the top candidate location(s) based on the average alignment scores of all supporting reads. It also implements stringent quality control procedures, i.e., reciprocal alignment of the supporting reads against the human reference sequence within each candidate region. For example, as a result of the customized BWA-MEM parameters, ERVcaller significantly increases the sensitivity for detecting novel and highly mutated TE insertions (e.g., when using the “LTR5_Hs” sequence as the only reference, ERVcaller was still capable of discovering insertions derived from related sequences, such as “LTR5A/B”).

The existing tools for the detection of TE insertions have significantly driven the past research into polymorphic TE insertions in the human genome. However, each of the existing tools still has some limitations. For example, TEMP, ITIS, and STEAK have no functions for genotyping. RetroSeq was able to genotype TE insertions; however, the genotyping function has been removed from its latest version. ITIS is not practical for analyzing large datasets due to computing time. Neither ITIS nor STEAK was able to efficiently detect partial ERV insertions (**Figure 3B**). MELT was newly designed to detect TE insertions; however, it only uses the aligned BAM files generated by BWA, limiting its utilization with BAM files from other aligners, and it only uses the reads with both ends unmapped within a one-million bp region, limiting the detection of the polymorphic TE insertions located close to fixed TEs. By comparison, ERVcaller was designed to efficiently detect and genotype TE insertions at single nucleotide resolution, regardless of the types and lengths of the TEs. ERVcaller is based on read alignment without the use of the prior reference repeat annotation information for detecting supporting reads or filtering false TE insertions, and thus it is likely to detect more TE insertions, regardless of TE locations. ERVcaller supports the use of both FASTQ and BAM files generated by BWA, Bowtie2 or other aligners. Not all published tools were compared in this study. For example, TIF (Nakagome, et al., 2014) was not analyzed because it only used split reads for TE detection. Tangram, which is also commonly used, was not included in this analysis due to installation issues (Rishishwar, et al., 2017; Wu, et al., 2014). **Supplementary Table 1** summarizes the comparison of the features of ERVcaller with the existing tools.

ERVcaller shares some of the limitations common to other NGS-based software. First, ERVcaller requires a TE reference(s) as it is based on sequence alignment. Second, it is challenging to identify the TE insertions that contain longer target site duplications at the breakpoints (Kahyo, et al., 2017). As a trade-off with high precision, ERVcaller would miss some of the TE insertions located in repetitive sequence regions, which is common to all tools of this kind. Third, due to the two LTRs of ERVs, it is challenging for all the tools, including ERVcaller, to determine their length, along with many other sequence features of identified ERV insertions using NGS reads with a short insert size, e.g., < 900 bp. These issues may be partially addressed by increasing insert size, read length, and depth of the NGS data, applying local *de novo* assembly, or using third-generation sequencing. Fourth, ERVcaller is designed to detect the TE insertions that are not present in the human reference genome. To detect the polymorphic TEs already present in the human reference genome, an alternate reference assembly may be created with all polymorphic TEs of interest removed. Alternatively, structural variation detection tools may be used to screen for polymorphic reference TEs as deletions.

ERVs have been reported to be associated with many human diseases. Many high frequency polymorphic ERV loci have been recently identified in human populations (Marchi, et al., 2014; Wildschutte, et al., 2016). Evidence suggests that there might be a large number of undiscovered polymorphic ERVs in the human genome (Moyes, et al., 2007). Indeed, in this study, ERVcaller newly identified eight polymorphic HERV-K loci in only 45 samples. Moreover, the number of polymorphic ERVs per individual and the frequencies of these ERVs vary significantly between different human samples (Wildschutte, et al., 2016). Presumably, some of the variable loci are associated with the human migration or susceptibility to certain diseases, and thus are population- or disease-specific. This suggests that polymorphic ERVs should be investigated to study their roles in human diseases and health, and to identify disease genes with the possibility of serving as genetic markers.

## Software availability

ERVcaller is an open-source software. ERVcaller v1.3 source code, documentation, and example data are available at www.uvm.edu/genomics/software/ERVcaller.html.

## Electronic database information

Accession numbers and URLs for data presented herein are as follows:

Repbase database: http://www.girinst.org;

dbRIP: http://dbrip.brocku.ca;

UCSC Genome Browser database: http://hgdownload.soe.ucsc.edu/downloads.html#human;

International Genome Sample Resource database: http://www.internationalgenome.org/home.

## Acknowledgement

The authors thank Sheryl White, and Thomas Buttolph in the COBRE Neuroscience Cellular and Molecular Core at the University of Vermont for their technical expertise in performing PCR and cloning. The authors thank the Vermont Integrative Genomics Resource for their technical expertise in performing Sanger sequencing. The authors thank the 1000 Genomes Project (Phase 3) and the colleagues for providing the sequencing data of their samples. The authors thank the Coriell Institute for Medical Research for providing DNA specimens. The authors thank the Vermont Advanced Computing Core and the Massachusetts Green High-Performance Computer C3DDB for providing computing resources. The authors thank Erik Brown for his assistance in the revision process of the manuscript. The authors thank Zhiping Weng, Akio Miyao, John Coffin, Arvis Sulovari, and others for their discussions and suggestions. The authors thank Jason Kost, Michael Mariani, Cong Gao, and John Baronas for their careful reviews of the manuscript. The authors thank the anonymous reviewers for their comments and suggestions.

## References

Bao, W., Kojima, K.K. and Kohany, O. Repbase Update, a database of repetitive elements in eukaryotic genomes. Mob DNA 2015;6(1):11.

Belshaw, R., et al. Genomewide screening reveals high levels of insertional polymorphism in the human endogenous retrovirus family HERV-K(HML2): implications for present-day activity. J Virol 2005;79(19):12507–12514.

Brodziak, A., et al. The role of human endogenous retroviruses in the pathogenesis of autoimmune diseases. Med Sci Monit 2012;18(6):RA80–88.

Burns, K.H. Transposable elements in cancer. Nat Rev Cancer 2017;17(7):415–424.

Chaisson, M.J.P., et al. Multi-platform discovery of haplotype-resolved structural variation in human genomes. bioRxiv 2018: 193144.

Chen, X., Kost, J. and Li, D. Comprehensive comparative analysis of methods and software for identifying viral integrations. Briefings in Bioinformatics 2018:bby070–bby070.

Chuong, E.B., Elde, N.C. and Feschotte, C. Regulatory evolution of innate immunity through co-option of endogenous retroviruses. Science 2016;351(6277):1083–1087.

Chuong, E.B., et al. Endogenous retroviruses function as species-specific enhancer elements in the placenta. Nat Genet 2013;45(3):325–329.

Danecek, P., et al. The variant call format and VCFtools. Bioinformatics 2011;27(15):2156–2158.

Douville, R.N. and Nath, A. Human endogenous retroviruses and the nervous system. Handb Clin Neurol 2014;123:465–485.

Fort, A., et al. Deep transcriptome profiling of mammalian stem cells supports a regulatory role for retrotransposons in pluripotency maintenance. Nat Genet 2014;46(6):558–566.

Fuentes, D.R., Swigut, T. and Wysocka, J. Systematic perturbation of retroviral LTRs reveals widespread long-range effects on human gene regulation. eLife 2018;7:e35989.

Gardner, E.J., et al. The Mobile Element Locator Tool (MELT): population-scale mobile element discovery and biology. Genome Res 2017;27(11):1916–1929.

Garrison, K.E., et al. T cell responses to human endogenous retroviruses in HIV-1 infection. PLoS Pathog 2007;3(11):e165.

Genomes Project, C., et al. A map of human genome variation from population-scale sequencing. Nature 2010;467(7319):1061–1073.

Goerner-Potvin, P. and Bourque, G. Computational tools to unmask transposable elements. Nature Reviews Genetics 2018;19(11):688–704.

Goke, J., et al. Dynamic transcription of distinct classes of endogenous retroviral elements marks specific populations of early human embryonic cells. Cell Stem Cell 2015;16(2):135–141.

Gonzalez-Cao, M., et al. Human endogenous retroviruses and cancer. Cancer Biol Med 2016;13(4):483–488.

Groger, V. and Cynis, H. Human Endogenous Retroviruses and Their Putative Role in the Development of Autoimmune Disorders Such as Multiple Sclerosis. Front Microbiol 2018;9:265.

Handsaker, R.E., et al. Discovery and genotyping of genome structural polymorphism by sequencing on a population scale. Nat Genet 2011;43(3):269–276.

Hormozdiari, F., et al. Combinatorial algorithms for structural variation detection in high-throughput sequenced genomes. Genome Res 2009;19(7):1270–1278.

Hormozdiari, F., et al. Next-generation VariationHunter: combinatorial algorithms for transposon insertion discovery. Bioinformatics 2010;26(12):i350–357.

Hu, X., et al. pIRS: Profile-based Illumina pair-end reads simulator. Bioinformatics 2012;28(11):1533–1535.

International Human Genome Sequencing, C. Initial sequencing and analysis of the human genome. Nature 2001;409:860.

Jern, P. and Coffin, J.M. Effects of retroviruses on host genome function. Annu Rev Genet 2008;42:709–732.

Jern, P. and Coffin, J.M. Effects of retroviruses on host genome function. Annu Rev Genet 2008;42(1):709–732.

Jiang, C., et al. ITIS, a bioinformatics tool for accurate identification of transposon insertion sites using next-generation sequencing data. BMC bioinformatics 2015;16(1):72.

Kahyo, T., et al. Insertionally polymorphic sites of human endogenous retrovirus-K (HML-2) with long target site duplications. BMC Genomics 2017;18(1):487.

Karamitros, T., et al. Human Endogenous Retrovirus-K HML-2 integration within *RASGRF2* is associated with intravenous drug abuse and modulates transcription in a cell-line model. Proceedings of the National Academy of Sciences 2018;115(41):10434–10439.

Kassiotis, G. Endogenous retroviruses and the development of cancer. Journal of immunology 2014;192(4):1343–1349.

Katzourakis, A., Pereira, V. and Tristem, M. Effects of Recombination Rate on Human Endogenous Retrovirus Fixation and Persistence. Journal of Virology 2007;81(19):10712–10717.

Kazazian, H.H. and Moran, J.V. Mobile DNA in Health and Disease. New England Journal of Medicine 2017;377(4):361–370.

Keane, T.M., Wong, K. and Adams, D.J. RetroSeq: transposable element discovery from next-generation sequencing data. Bioinformatics 2013;29(3):389–390.

Lee, E., et al. Landscape of somatic retrotransposition in human cancers. Science 2012;337(6097):967–971.

Leung, A., et al. LTRs activated by Epstein-Barr virus-induced transformation of B cells alter the transcriptome. Genome Research 2018.

Li, H. Aligning sequence reads, clone sequences and assembly contigs with BWA-MEM. *arXiv preprint arXiv:1303.3997* 2013.

Li, H., et al. The Sequence Alignment/Map format and SAMtools. Bioinformatics 2009;25(16):2078–2079.

Li, W., et al. Human endogenous retrovirus-K contributes to motor neuron disease. Sci Transl Med 2015;7(307):307ra153.

Macfarlane, C. and Simmonds, P. Allelic variation of HERV-K(HML-2) endogenous retroviral elements in human populations. J Mol Evol 2004;59(5):642–656.

Macfarlane, C.M. and Badge, R.M. Genome-wide amplification of proviral sequences reveals new polymorphic HERV-K(HML-2) proviruses in humans and chimpanzees that are absent from genome assemblies. Retrovirology 2015;12(1):35.

Marchi, E., et al. Unfixed endogenous retroviral insertions in the human population. J Virol 2014;88(17):9529–9537.

Marguerat, S., et al. Association of human endogenous retrovirus K-18 polymorphisms with type 1 diabetes. Diabetes 2004;53(3):852–854.

Mills, R.E., et al. Which transposable elements are active in the human genome? Trends Genet 2007;23(4):183–191.

Moyes, D., Griffiths, D.J. and Venables, P.J. Insertional polymorphisms: a new lease of life for endogenous retroviruses in human disease. Trends Genet 2007;23(7):326–333.

Nakagome, M., et al. Transposon Insertion Finder (TIF): a novel program for detection of de novo transpositions of transposable elements. BMC bioinformatics 2014;15(1):71.

Navarro, C. The Mobile World of Transposable Elements. Trends in Genetics 2017;33(11):771–772.

Quinlan, A.R., et al. Genome-wide mapping and assembly of structural variant breakpoints in the mouse genome. Genome Res 2010;20(5):623–635.

Rishishwar, L., Mariño-Ramírez, L. and Jordan, I.K. Benchmarking computational tools for polymorphic transposable element detection. Briefings in Bioinformatics 2017;18(6):908–918.

Robbez-Masson, L. and Rowe, H.M. Retrotransposons shape species-specific embryonic stem cell gene expression. Retrovirology 2015;12:45.

Rooney, M.S., et al. Molecular and genetic properties of tumors associated with local immune cytolytic activity. Cell 2015;160(1–2):48–61.

Santander, C.G., et al. STEAK: A specific tool for transposable elements and retrovirus detection in high-throughput sequencing data. Virus Evol 2017;3(2):vex023.

Slokar, G. and Hasler, G. Human Endogenous Retroviruses as Pathogenic Factors in the Development of Schizophrenia. Front Psychiatry 2015;6:183.

Stewart, C., et al. A comprehensive map of mobile element insertion polymorphisms in humans. PLoS Genet 2011;7(8):e1002236.

Sudmant, P.H., et al. An integrated map of structural variation in 2,504 human genomes. Nature 2015;526(7571):75–81.

The Genomes Project, C. A global reference for human genetic variation. Nature 2015;526(7571):68–74.

Thomas, J., Perron, H. and Feschotte, C. Diverse endogenous retroviruses generate structural variation between human genomes via LTR recombination. *bioRxiv* 2018.

Thung, D.T., et al. Mobster: accurate detection of mobile element insertions in next generation sequencing data. Genome biology 2014;15(10):488.

Untergasser, A., et al. Primer3—new capabilities and interfaces. Nucleic Acids Research 2012;40(15):e115–e115.

Wang, J., et al. dbRIP: a highly integrated database of retrotransposon insertion polymorphisms in humans. Hum Mutat 2006;27(4):323–329.

Wessler, S.R. Transposable elements and the evolution of eukaryotic genomes. Proceedings of the National Academy of Sciences 2006;103(47):17600–17601.

Wildschutte, J.H., et al. Discovery of unfixed endogenous retrovirus insertions in diverse human populations. Proceedings of the National Academy of Sciences of the United States of America 2016;113(16):E2326–2334.

Wu, J., et al. Tangram: a comprehensive toolbox for mobile element insertion detection. BMC Genomics 2014;15(1):795.

Zhuang, J., et al. TEMP: a computational method for analyzing transposable element polymorphism in populations. Nucleic Acids Res 2014;42(11):6826–6838.

